# Single-cell mechanical analysis reveals viscoelastic similarities between normal and neoplastic brain cells

**DOI:** 10.1101/2023.09.23.559055

**Authors:** Killian Onwudiwe, Julian Najera, Luke Holen, Alice A. Burchett, Dorielis Rodriguez, Maksym Zarodniuk, Saeed Siri, Meenal Datta

## Abstract

Understanding cancer cell mechanics allows for the identification of novel disease mechanisms, diagnostic biomarkers, and targeted therapies. In this study, we utilized our previously established fluid shear stress assay to investigate and compare the viscoelastic properties of normal immortalized human astrocytes (IHAs) and invasive human glioblastoma (GBM) cells when subjected to physiological levels of shear stress that are present in the brain microenvironment. We used a parallel-flow microfluidic shear system and a camera-coupled optical microscope to expose single cells to fluid shear stress and monitor the resulting deformation in real-time, respectively. From the video-rate imaging, we fed cell deformation information from digital image correlation into a three-parameter generalized Maxwell model to quantify the nuclear and cytoplasmic viscoelastic properties of single cells. We further quantified actin cytoskeleton density and alignment in IHAs and GBM cells via immunofluorescence microscopy and image analysis techniques. Results from our study show that contrary to the behavior of many extracranial cells, normal and cancerous brain cells do not exhibit significant differences in their viscoelastic behavior. Moreover, we also found that the viscoelastic properties of the nucleus and cytoplasm as well as the actin cytoskeletal densities of both brain cell types are similar. Our work suggests that malignant GBM cells exhibit unique mechanical behaviors not seen in other cancer cell types. These results warrant future study to elucidate the distinct biophysical characteristics of the brain and reveal novel mechanical attributes of GBM and other primary brain tumors.

## 1. Introduction

Tissue microenvironments of different organs possess distinct biological and mechanical features that uniquely influence the biophysical responses and characteristics of cancer cells [1-8]. Notably, in organs such as the breast, ovary [9], thyroid [10], bladder [11], kidney [12], prostate [13], and lung, significant differences in the viscoelastic properties of cancer cells relative to their normal cellular counterparts are used as indicators of cancer onset and/or progression [14-18]. Cancer cells are typically softer (i.e., less stiff) and less viscous than normal cells, allowing them to deform and migrate freely within the tumor, away from the tumor, and to distant sites. While these mechanical distinctions have been well documented for the tissue sites [14-20], it is unknown if these cellular mechanical differences exist in the case of glioblastoma (GBM), an aggressive primary brain cancer. In this study, we compared the nuclear and cytoplasmic viscoelastic behavior of the human GBM cell line U87 to normal immortalized human astrocytes (IHAs), the purported cell of origin of GBM. Using our previously established microfluidic device [21], we applied physiological levels of fluid shear stress found in the brain environment [22] to adherent GBM cells and IHAs and imaged their deformation in real-time to compute stiffness, viscosity, and relaxation time on a single-cell basis via downstream digital image correlation (DIC) and mechanical modeling. We also quantified actin abundance, organization, and localization as potential cytoskeletal determinants of the observed mechanical behaviors. We found that the viscoelastic properties of GBM cells were strikingly similar to those of IHAs – a stark contrast from the mechanical disparities previously found between normal and neoplastic cells of other organ sites. Furthermore, IHAs maintain viscoelastic differences between the nucleus and cytoplasm, but these cellular compartments are not mechanically distinct in GBM cells. Finally, while total cytoskeletal content was found to be similar between normal and cancerous brain cells, IHAs had more aligned actin fibers than GBM cells.

## 2. Materials and Methods

### 2.1. Cell Culture

IHAs (Creative Bioarray, CSC-C12025Z, Shirley, NY, USA) and U87 cells (ATCC, Manassas, VA, USA) were cultured at 37 °C in a humidified 5% CO_2_ atmosphere. GBM cells were cultured in Dulbecco’s Modified Eagle Medium (Invitrogen, Carlsbad, CA, USA) supplemented with 10% fetal bovine serum (Gibco, Grand Island, NY, USA) and 1% Penicillin-Streptomycin (Invitrogen, Carlsbad, CA, USA), and IHAs were cultured in SuperCult IHA Media (Creative Bioarray, Shirley, NY, USA), 1% IHA growth supplement (Creative Bioarray, Shirley, NY, USA), 2% fetal bovine serum (Creative Bioarray, Shirley, NY, USA) and 1% Penicillin-Streptomycin (Creative Bioarray, Shirley, NY, USA). All cells were cultured in 35 mm Falcon dishes (Corning Inc., Corning, NY, USA) for 48 h to ensure proper attachment and spreading on the substrate. Cells were seeded at a density of 5,000 cells per petri dish to ensure that single cells could be identified and so that single-cell deformation was not significantly affected by neighboring cells.

### 2.2. Shear Assay

The mechanical properties of IHAs and U87 cells were evaluated using our previously established modified shear assay technique. A schematic of the system and the visualized process can be found in our recent publication [21]. For a more viscous shear fluid media, 0.5 wt% non-allergenic and non-toxic methylcellulose (Sigma Aldrich, St. Louis, MO, USA) was added to the serum-free DMEM culture media. The viscosity of the media (0.017Pa. s) was determined using an HR-20 rheometer (TA Instruments, 159 Lukens Drive, New Castle, DE 19720). The applied wall shear stress was calculated using the following formula for a parallel-plate flow chamber:

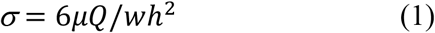

where *μ* is the viscosity of the fluid, Q is the volumetric flow rate, and *w* and *h* are the width and height of the rubber gasket, respectively. Using our shear assay apparatus, we subjected cells to a shear stress of 4 Pa, the physiological level of shear stress experienced by cells in the brain [22]. Phase-contrast video recordings (Nikon Microscope, Brighton, MI, USA) were taken of cells while exposed to shear stress for 8 minutes. A total of 84 U87 cells and 96 IHAs were analyzed for mechanical characterization using DIC analysis.

### 2.3. Digital Image Correlation

DIC was performed using parameters described in previous work and visualized experiments publications [14, 16, 21]. Deformation and strain within the cell are tracked by taking the sum of differential of the deformation of an image with respect to the previous images (Fig 1A – E). For a given cell image, we used a subset size of 31x31 pixels and a step size (i.e., deformation distance on each subset) of 20 pixels. Nuclear and cytoplasmic deformation and strain behavior are determined by analyzing different representative areas of the nucleus and cytoplasm of the cells using strain gauges (Fig 1F – I). An average of 3 points (area of interest) were selected both in the nucleus and cytoplasm of each cell. The output from the correlation analysis is strain-time data that can be further analyzed using MATLAB to determine the mechanical properties of each cell. A more detailed step-by-step protocol can be found in our recently published article [21].

**Figure 1.**
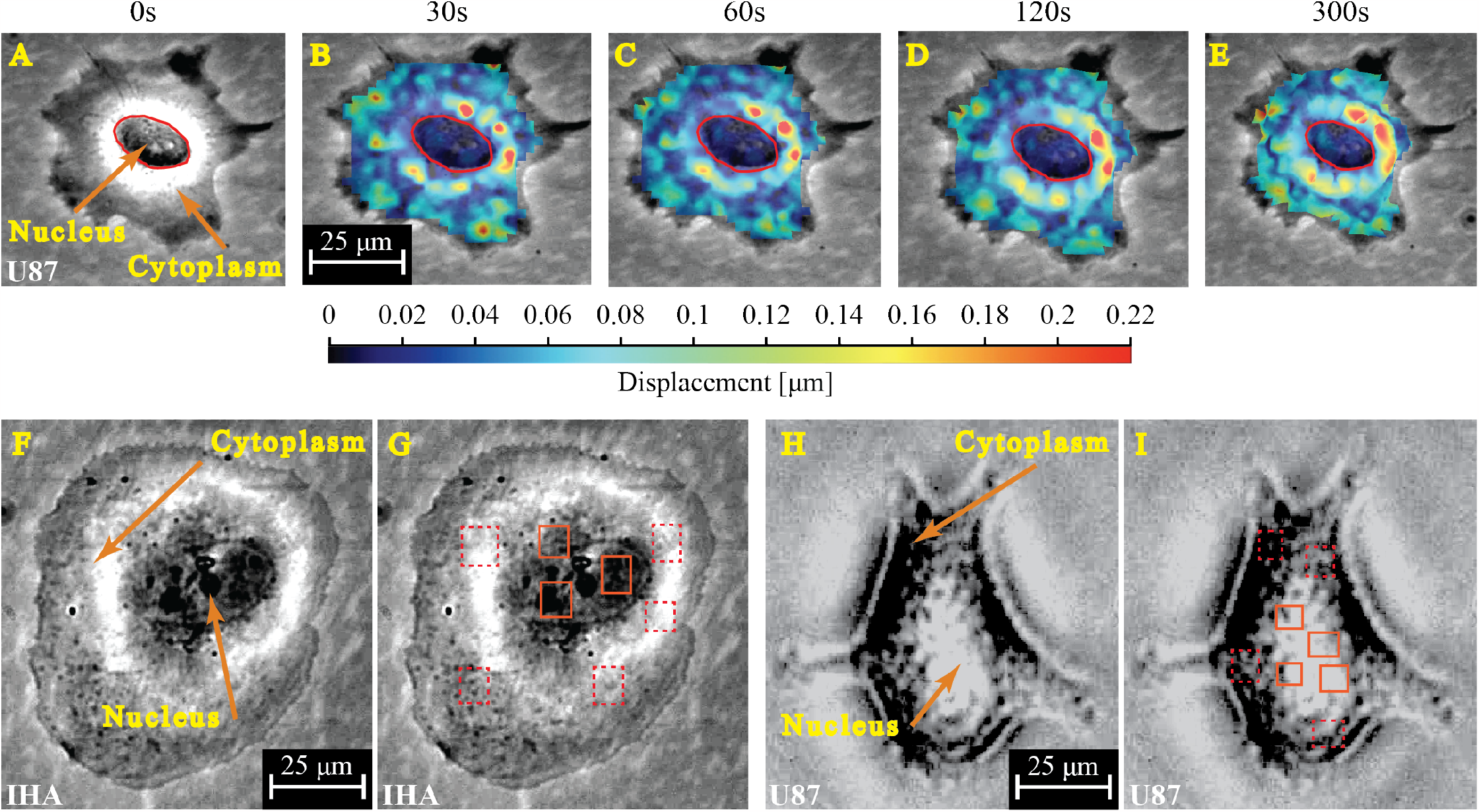
DIC reveals varying time- and spatially-dependent displacement and strain magnitudes across a single cell under shear stress. In this representative image of a GBM cell (A–E), cytoplasmic regions exterior to the nucleus (red outline) exhibit heterogeneous mechanical responses with higher displacement and strain rates compared to the nucleus. (A) Cell before shear stress application. (B–E) Evolution of displacement and strain with time. (F–I) Phase-contrast images and DIC pixel block placement for the analysis of the nucleus and cytoplasm of IHAs (F&G) and GBMs (H&I).

### 2.4. Fluorescent Staining and Imaging

To assess cytoskeletal organization, fluorescent staining was performed on IHAs and U87 GBM cells in line with our previously reported experimental procedures. The stained cells were imaged using a Keyence BZ-X810 widefield microscope (Keyence, Itasca, IL, USA).

### 2.5. Actin Density Quantification and Fiber Alignment Analysis

The Leica fluorescence image processing software LAS X (Leica Microsystems, IL, USA) was used to quantify individual cellular actin density for the two cell lines. Single cells were analyzed by mapping out a region of interest (ROI) that corresponds to the area of the cell. The mean actin stain intensity and the area of the ROI in each channel were quantified by the software, resulting in a mean intensity/area (a.u./μm^2^) for approximately 100 of each cell type. For fiber alignment analysis, widefield fluorescent images were cropped to contain single cells, and intensity was scaled so that the maximum intensity value for each was set to 255. These images were then processed by the MATLAB-based software CT-FIRE to identify fibers and their corresponding angle values. The angles were normalized to the circular mean angle for each cell and combined for statistical analysis. The analyzed image set contained 32 IHAs and 29 U87 cells.

### 2.6. Statistical Analysis

Statistical tests (t-tests) were performed using the IBM SPSS statistical software (Chicago, IL, USA). A confidence level of 95% was adopted and statistical differences between the means were considered significant at *p* < 0.05. Bar plots display the mean ± standard error of the mean (SEM). Cytoskeletal fiber alignment was analyzed in MATLAB using the Circular Statistics toolbox, the Kruskal-Wallis function, and the Kolmogorov-Smirnov function.

## 3. Modeling and Computation

### 3.1. Viscoelastic Modeling

In this study, we employed a three-element generalized Maxwell model to characterize the viscoelastic properties of cells, as described in our previous work [21]. The model’s constitutive equation describes the relationship between stress and strain by combining a dashpot and spring in series. The dashpot represents the viscous behavior of the cellular material, dissipating energy as the cell undergoes deformation over time. This accounts for the time-dependent nature of cellular responses, such as stress relaxation. Meanwhile, the ring component captures the elastic response, storing energy as the cell is deformed and restoring the original shape after the force is removed. By incorporating both elements, the model provides a comprehensive representation of the viscoelastic behavior observed in biological cells. The constitutive equation guiding this principle is shown in equations 2 & 3:

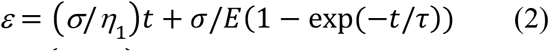

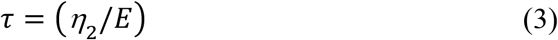

where ε represents the strain, 3 represents the stress, E represents the stiffness, *η*_1_ represents the effective cellular viscosity.*η*_2_ represents the secondary dashpot viscosity from which the relaxation time, τ, is obtained. This equation was used to fit the strain-time data obtained from the shear experiment and extracted by the DIC software. The curve fitter tool in MATLAB and the custom equation option were used to fit the data and execute the model. Hence the model successfully characterized and generated the corresponding viscoelastic properties (stiffness, viscosity, and relaxation time).

## 4. Results

### 4.1. IHAs and GBM Cells Share Deformation and Creep Responses Under Shear Stress

The strain-time behavior of cells reflects their ability to withstand deformation under applied stress and reveals characteristic creep deformation regimes that are significant for viscoelastic material responses. The primary (elastic), secondary (steady state), and tertiary (failure) regimes, represent the different phases of viscoelastic deformation. Fig. 2 shows the deformation behavior of IHAs and GBM cells when subjected to 4 Pa of fluid shear stress for 8 minutes. Our analysis shows that IHAs and GBMs experience similar creep deformation behavior. However, these responses are distinct between the nucleus and cytoplasm of both cells. We find that the cytoplasm of both cells experiences slightly higher deformation and strain compared to the nucleus (Fig. 2-F).

**Figure 2.**
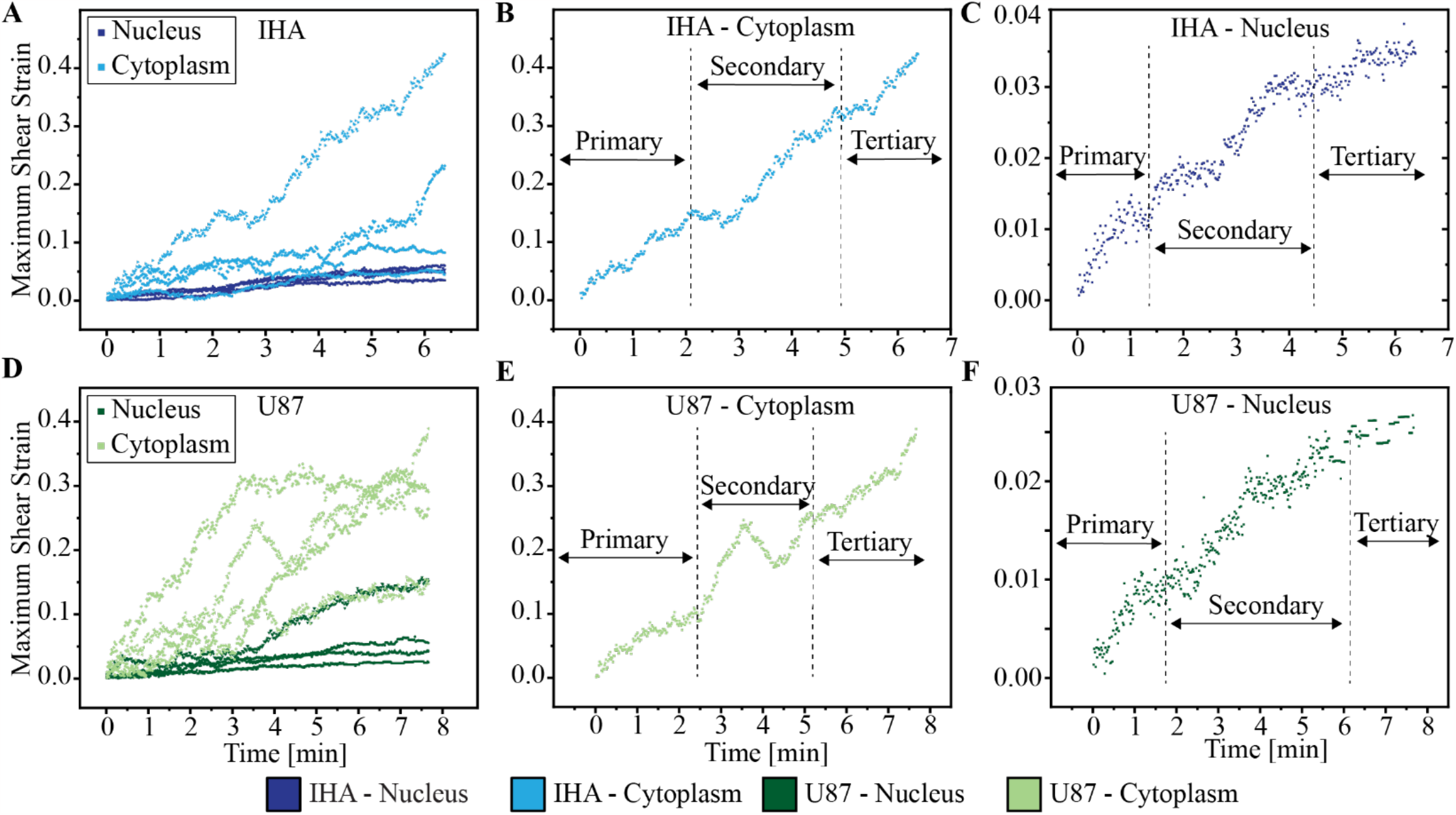
IHAs and GBM cells have similar strain-time deformation profiles. Representative strain-deformation profiles of the nucleus and cytoplasm of IHAs (A) and GBM cells (B) divided into primary, secondary, and tertiary regimes which correspond to elastic, steady-state, and plastic/failure phases during deformation, respectively (C-F). Their characteristic responses at each phase can be explored to determine their viscoelastic properties in response to applied stress.

### 4.2. IHAs and GBM Cells Exhibit Similar Viscoelastic Properties

We next compared the mechanical properties of the IHAs to those of GBM cells. Surprisingly, IHAs and GBM cells exhibit similar viscoelastic properties. As seen in Fig. 3, there are no distinguishable mechanical differences in the nuclei or cytoplasm between the normal and malignant brain cells. Specifically, differences between the stiffness, viscosity, and relaxation times of IHAs and GBM cells are statistically insignificant (Fig. 3). This unique similarity between healthy and cancerous brain cells may be a relationship that distinguishes glioblastoma from other cancer types and could highlight a distinct mechanical cancer-host interaction than what is found elsewhere.

**Figure 3:**
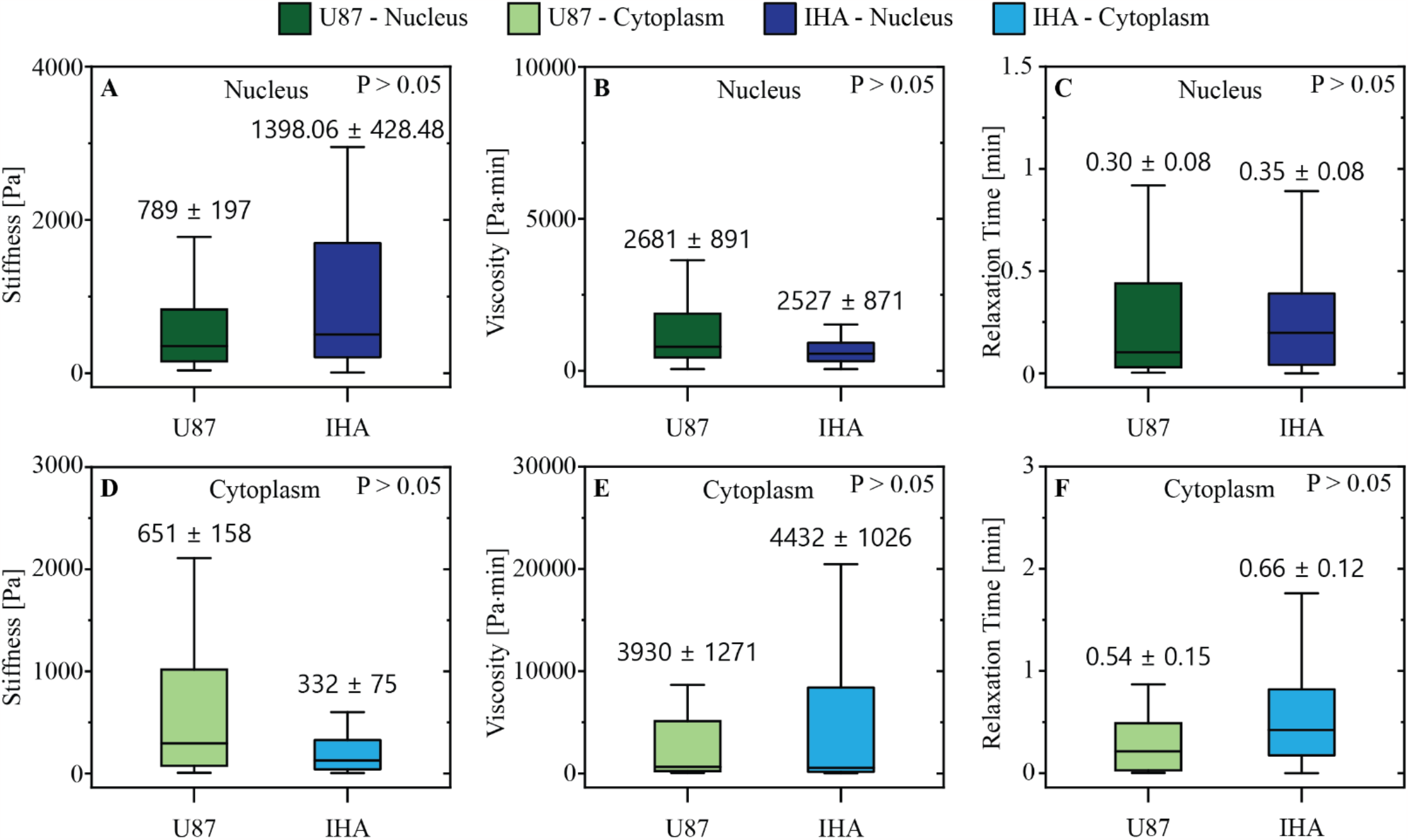
GBM cells and IHAs have similar viscoelastic properties. Under shear stress, (A) nuclear stiffness, (B) nuclear viscosity, (C) nuclear relaxation time, (D) cytoplasmic stiffness, (E) cytoplasmic viscosity, and (F) cytoplasmic relaxation time do not differ between IHAs and GBM cells (all p-values > 0.05).

### 4.3. IHAs – but not GBM cells – Feature Intracellular Compartmental Variations Between Nuclear and Cytoplasmic Viscoelasticity

Previous studies have established that the nucleus is stiffer than the surrounding cytoplasm in many cell types [16, 24, 25]. To determine whether these differences exist between normal IHAs and their malignant counterpart, we characterized the viscoelastic properties of their nuclear and cytoplasmic regions using the three-element generalized Maxwell model. Our results show that distinct nuclear and cytoplasmic viscoelastic behaviors exist in the IHAs, but not in the GBM cells. In particular, the average nuclear stiffness of IHAs (1398 ± 428 Pa) is four-fold higher than the stiffness of the cytoplasm (331 ± 74 Pa) (Fig. 4A). Similarly, the nuclear relaxation time of IHAs (0.35 ± 0.08 min) (Fig. 4C) is significantly lower than the cytoplasmic relaxation time (0.66 ± 0.12 min). However, there is no statistical difference between the viscosity of the nucleus (2526 ± 870 Pa•min) and cytoplasm (4432 ± 1026.35 Pa•min) in IHAs (Fig. 4B). In comparison, none of the viscoelastic properties of GBM cells differ significantly between the nucleus and cytoplasm (Fig. 4D-F).

**Figure 4.**
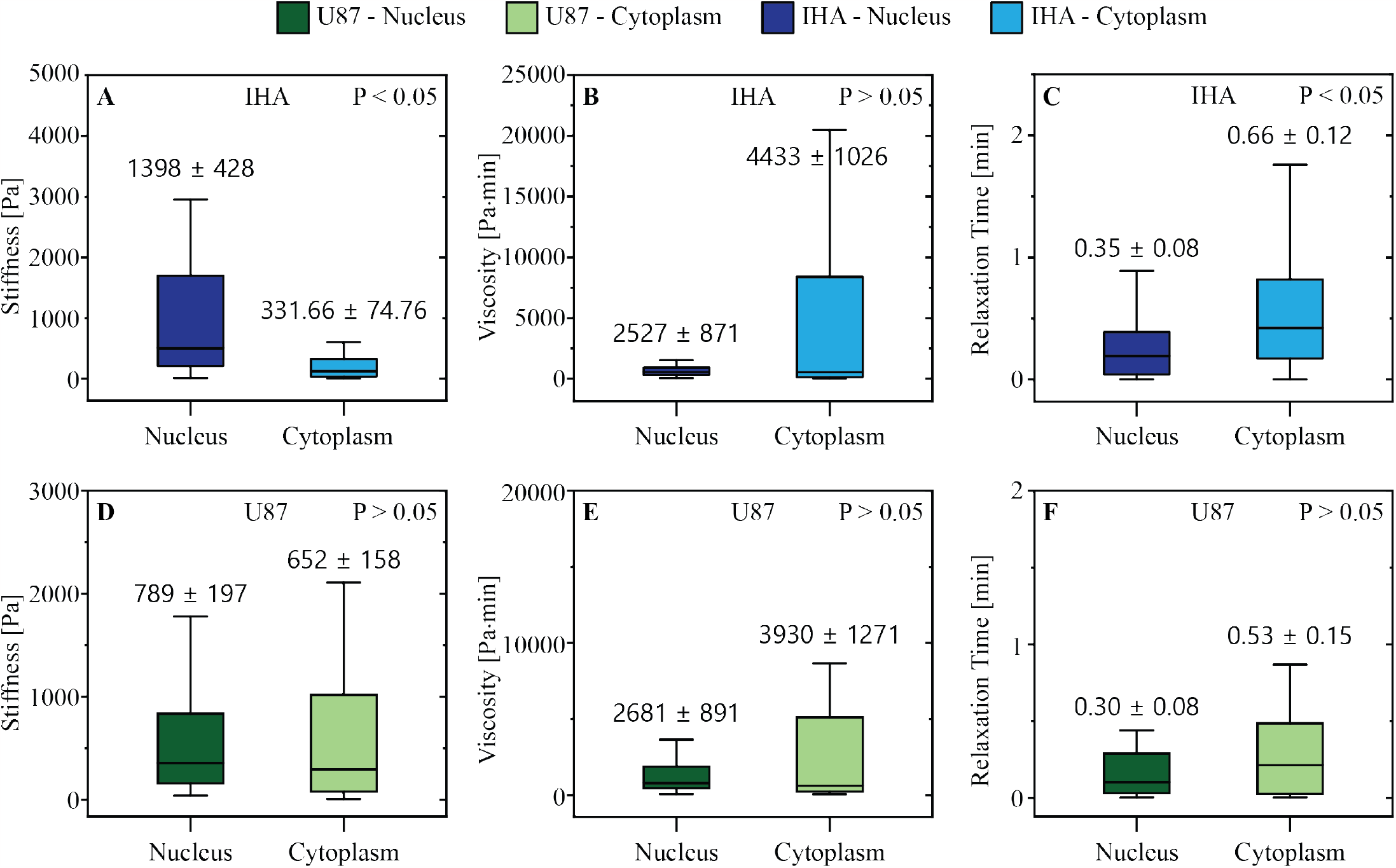
IHA intracellular compartments feature viscoelastic differences that are not conserved in GBM cells. Comparison between IHA nuclear and cytoplasmic (A) stiffness, (B) viscosity, and (C) relaxation time reveals notable differences in intracellular stiffness and relaxation time. However, GBM cells do not have significant differences in cytoplasmic versus nuclear (D) stiffness, (E) viscosity, and (F) relaxation time.

### 4.4. IHAs and GBM cells Share Similar Actin Content but Different Alignment

We next used fluorescence staining to investigate actin cytoskeletal content and alignment as this protein is known to regulate and govern cell mechanics and behavior [26-28]. The average actin stain intensity per unit area is similar between the IHAs and GBM cells (Fig. 5A-C). However, the degree of actin fiber organization differs between the two cell types. Fibers in individual IHAs are more likely to be aligned in a single mean direction, whereas the fibers of GBM cells have a more random angle distribution that is significantly different from the IHAs (Fig. 5D-E). Specifically, GBM cells have increased variance in angle orientation within each cell (p = 0.0019) and a decrease in the angle distribution kurtosis (p = 0.00043) compared to IHAs, thus reflecting the more random distribution of fiber angles in GBM cells. This distinction between the two cell types may have a physiological significance that should be further explored in future work.

**Figure 5.**
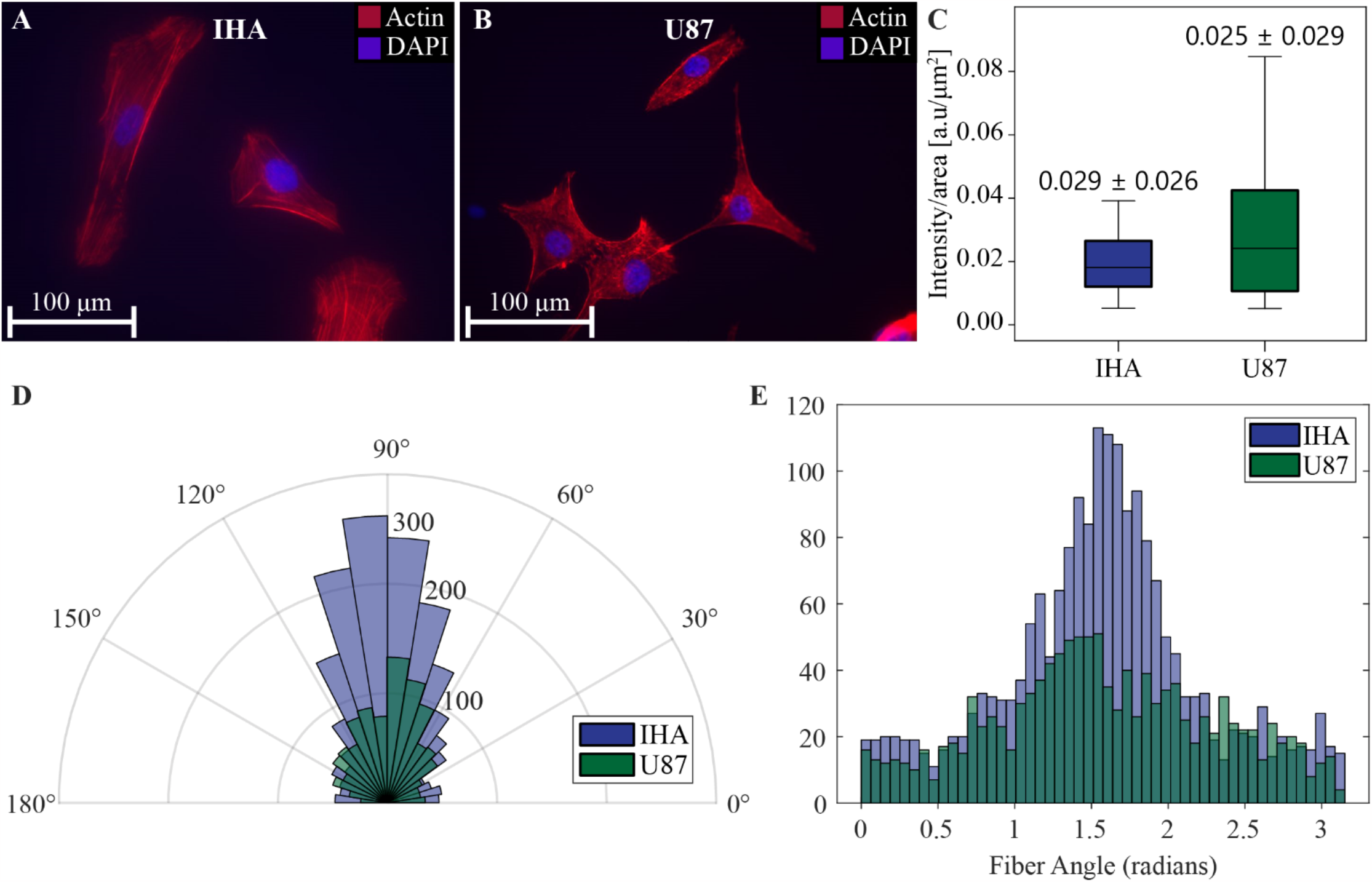
GBM cells and IHAs have similar actin content but distinct actin organization and localization. Representative phalloidin-stained (A) GBM cells and (B) IHAs. (C) The total cell average actin stain intensity is not significantly different between GBM cells and IHAs. (D) Angular histogram representation of fiber orientation in GBM cells and IHAs, centered to 90 degrees for each cell. (E) The distribution of fiber angles relative to the average angle in each cell is distinct, with IHA fibers being more aligned than GBM fibers, centered to 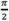 radians.

### 4.5. Distinct Variations in the Mechanical Properties of Cancer and Normal Cells Exist in Several Organs but not in the Brain

To highlight the unique viscoelastic behavior of brain cells, we compared our results to previous studies that performed a similar mechanical characterization technique (i.e., fluid shear stress assay) on other cell types (Table 1). We find that significant viscoelastic differences exist between cancerous cells and their healthy cellular counterparts in other organ systems (e.g., particularly in the case of breast cells), which is in contrast to our results with brain cells.

**Table 1.** Viscoelastic Properties of Cancerous Cells and their Normal Counterparts Across Different Organs Using Shear Assay Technique.

| Tissue of Origin | Cell | Cell state | Nuclei Stiffness [Pa] | Cytoplasm Stiffness [Pa] | Nuclei Viscosity [Pa-min] | Cytoplasm Viscosity [Pa-min] | Ref. |
| --- | --- | --- | --- | --- | --- | --- | --- |
| Brain | IHA | Normal | 1398 | 332 | 2527 | 4432 | Current study |
|  | *U87 | Cancer | 789 | 652 | 2681 | 3930 |  |
| Breast | MCF-10A | Normal | 7966 | 3598 | 6868 | 3132 | [16] |
|  | *MDA-MB-468 | Cancer (Less Metastatic) | 5632 | 3334 | 5616 | 3390 |  |
|  | **MDA-MB-231 | Cancer (Highly Metastatic) | 622 | 261 | 325 | 215 |  |
| Bone | HOS | Cancer | 2740 | 715 | 6353 | 2138 | [15] |

\*\* Significant difference between cancer cells and normal cells
\*No significant difference between cancer cells and normal cells

## 5. Discussion

Mechanical responses and viscoelastic properties of normal and neoplastic brain cells are presented here in the context of physiological fluid shear stresses found in the brain [22]. We first show that IHAs and GBM cells exhibit similar creep behavior in response to applied stress, which is characteristic of the time-dependent deformation of viscoelastic materials under a constant load. For both IHAs and GBM cells, the cytoplasm experiences more creep compared to the nucleus, which is more compact and homogenous.

Distinct intracellular regions (such as the nucleus and cytoplasm) have been shown to display varying viscoelastic properties in other cell types [16]. Our findings show that the nucleus and cytoplasm of IHAs have distinct stiffness and relaxation times. However, GBM cells do not appear to have significant differences in their nuclear and cytoplasmic viscoelastic properties. Moreover, when comparing the viscoelastic properties of both cell types with one another, there are no significant mechanical differences between IHAs and GBM cells – a finding that appears to be unique to the brain. Further exploration of other experimental parameters, such as the mechanical properties of extracellular matrix and/or cell culture dish substrate coatings, may reveal other mechanical differences not captured in this study.

The actin cytoskeleton plays a pivotal role in governing cellular mechanics [29-31]. In this study, we found no significant differences in actin cytoskeleton abundance between the IHAs and GBM cells. These results align with our observation that there are no statistically significant differences in the viscoelastic properties (stiffness, viscosity, and stress relaxation) between these two cell types. However, IHAs have a more organized and aligned actin cytoskeleton compared to GBM cells, which may contribute to subtle differences in mechanical and/or physiological behavior that are perhaps not captured in the current study; this warrants further investigation.

Comparing the relationship between the viscoelastic properties of the brain and cancer cells investigated in this study with those of other organs [12, 15, 16, 32-38] suggests that the behavior of cancer cells in the brain (e.g., migration, motility, metabolism, proliferation, and metastasis) diverges considerably from those in extracranial organs [39]. The brain, an organ whose primary function does not involve mechanical load bearing (e.g., as in bone or muscle), is comparatively soft and may provide a less mechanically hostile microenvironment. Given the essential roles of cellular stiffness and viscosity in cancer cell migration [40], along with the low mechanical demands and high fluid content in the brain, cancer cells may adapt brain-specific mechanical properties via inherent (genetic) and externally-driven (microenvironmental) mechanisms. This may in part explain why GBM rarely seeds secondary nodes within the brain away from the primary site nor metastasizes to extracranial sites, although further studies are needed to connect these phenomena.

## 6. Conclusion

We explored the viscoelastic properties of normal and malignant brain cells as well as their cytoskeletal determinants. Our findings indicate that GBM cells – unlike cancer cells in other regions of the body – are mechanically similar to normal brain cells, although they lack the intracellular compartment-specific differences seen in IHAs. The unique mechanical properties of primary brain cancer cells may in part mediate their preferential local growth at their site of origin in the brain, rather than spreading to other parts of the cerebrum or body. Further research is required to understand the genetic and microenvironmental factors that modulate the mechanical behavior of healthy and cancerous cells in the brain.

## Acknowledgments

We thank Ms. R’nld Rumbach for her technical support. We thank Notre Dame Research for providing resources to support our initial investigations of IHAs. We thank the Notre Dame Chemical and Biomolecular Engineering Department for supporting and organizing Ms. Rodriguez’s research experience for undergraduates (REU). Fluorescent imaging was carried out in the Notre Dame Integrated Imaging Facility at the University of Notre Dame, with the help of Dr. Sara Cole. We thank Dr. Robert Nerenberg, Ms. Yanina Nahum, and the Materials Characterization Facility for their input and assistance with rheometric studies. We thank Drs. Holly Goodson and Christopher Patzke for providing staining antibodies.

